# The circadian clock mediates daily bursts of cell differentiation by periodically restricting cell differentiation commitment

**DOI:** 10.1101/2022.05.17.492365

**Authors:** Zhi-Bo Zhang, Joydeb Sinha, Zahra Bahrami-Nejad, Mary N. Teruel

## Abstract

Most mammalian cells have an intrinsic circadian clock that coordinates their metabolic activity with the daily rest and wake cycle. In addition, the circadian clock is known to regulate cell differentiation, but how continuous daily oscillations of the internal clock control a much longer, multi-day differentiation process is not known. Here we simultaneously monitor the circadian clock and progression of adipocyte differentiation live in single cells. Strikingly, we find a bursting behavior in the cell population whereby individual preadipocytes commit to differentiate primarily during a 12-hour window each day corresponding to the time of rest. Daily gating of differentiation occurs because cells can irreversibly commit to differentiate within a few hours, which is faster than the rest phase and much faster than the overall multi-day differentiation process. We show that the daily bursts in differentiation are driven by a variable and slow increase in expression of PPARG, the master regulator of adipogenesis, combined with rapid, clock-driven expression of CEBPA, which is in a fast positive feedback relationship with PPARG. During each rest cycle, the increase in CEBPA causes a brief step increase in PPARG so that some cells can reach the threshold to irreversibly commit to differentiate, causing the consecutive daily bursts in cell differentiation at the population level. Our findings are broadly relevant given that most differentiating somatic cells are regulated by the circadian clock. Having a restricted time each day when differentiation occurs may open therapeutic strategies to use timed treatment relative to the clock to promote tissue regeneration.

**Significance Statement:** Cells rely on a circadian clock that coordinates cellular activities with the day-night cycle. Defects in circadian clock genes dysregulate cell differentiation processes in immune, muscle, skin and fat cells. However, how a perpetual daily clock can regulate a multi-day long cell differentiation process was not understood. Here we show that the circadian clock controls a fast upregulation of the transcription factor CEBPA during each daily rest phase which in turn controls a fast irreversible step during the overall slow multi-day differentiation of fat cells, causing daily bursts of cell differentiation. Our finding opens potential therapeutic strategies to enhance tissue regeneration by timing when during the day drugs are administered.

## Introduction

Virtually all cells in the human body contain an intrinsic circadian clock (cell-intrinsic clock), operated by a set of core clock proteins that engage in coupled positive and negative transcriptional and translational feedback loops to generate rhythmic expression of 10-15% of the transcriptome(1). When components that drive the cell-intrinsic clock are genetically perturbed, cell differentiation of fat cells (adipocytes)(2, 3), T-cells(4), myoblasts(5), and embryonic stem cells(6) are defective, suggesting that the circadian clock regulates differentiation. However, it is not clear how a daily clock that oscillates perpetually can control a much slower process such as cell differentiation, which typically takes several days or even weeks.

One possibility is that cells count the number of circadian cycles to delay differentiation for a certain time period after the differentiation stimulus is added. Another possibility is that there may be a time window during each circadian cycle in which cells have a fast step during the overall slow differentiation process, which may prolong differentiation if a cell misses to differentiate and needs to wait for a subsequent permissive window. To distinguish between these and other possible mechanisms, we used adipogenesis as a model system since it is currently the only differentiation system for which validated tools are available to measure in live cells the time when cells irreversibly differentiate. Our strategy builds on a previously developed method to track cell differentiation progression by monitoring the endogenous expression of PPARG, the master regulator of adipogenesis, over several days(7, 8). The time when a preadipocyte irreversibly differentiates, called the differentiation commitment point, can be measured as the time when the PPARG level in the cell increases to a critical threshold at which positive feedback loops engage to lock the PPARG level at a perpetually high level(7, 8). Here we sought to understand how the circadian clock controls differentiation by using this cell model to measure the circadian clock and differentiation commitment live in the same cell.

Strikingly, rather than finding evidence for a counting of circadian cycles, we found that preadipocytes pass the commitment point in repeated daily bursts that occur exclusively during the phase of the circadian cycle which matches the resting period in humans(9). Mechanistically, we show that circadian expression of CEBPA, a positive feedback regulator of PPARG, controls a periodic increase in PPARG which then triggers differentiation only if PPARG reaches the threshold during the resting phase of the circadian cycle. Even though differentiation takes many days, a fast, only hour-long kinetic step, which increases PPARG from a lower to a higher intermediate level, drives irreversibly commitment, explaining how circadian rhythms can control a differentiation process that takes many days. Our study argues that the cell-intrinsic circadian clock controls cell differentiation by restricting it to a short phase window each day, providing a mechanism for how dysregulated circadian rhythms may broaden this daily phase window to increase differentiation and fat mass.

## Results

### Development of a system to simultaneously monitor the cell-intrinsic clock and cell differentiation progression in single cells

Adipogenesis is a multi-day process during which preadipocytes irreversibly differentiate into adipocytes primarily through the expression of PPARG, the master regulator of fat cell differentiation (Fig. 1a). Our previous studies showed that the time when preadipocytes irreversibly commit to becoming adipocytes, also known as the adipogenesis commitment point, can be precisely marked by the time at which the abundance of PPARG protein reaches a threshold level(7, 8). To understand how the cell-intrinsic clock regulates the timing of adipogenesis, we used a modified version of a previously described circadian reporter(10), which is comprised of coding and promoter sequences of *Rev-Erb*a conjugated to mScarlet (RFP) protein. We introduced this *Rev-Erb*a circadian clock reporter into an OP9 preadipocyte cell line in which endogenous PPARG had been tagged with citrine (YFP) using CRISPR genome editing(7). Figure 1b shows example time courses of citrine-PPARG / *Rev-Erb*a-mScarlet dual reporter cells undergoing adipogenesis.

**Figure 1.**
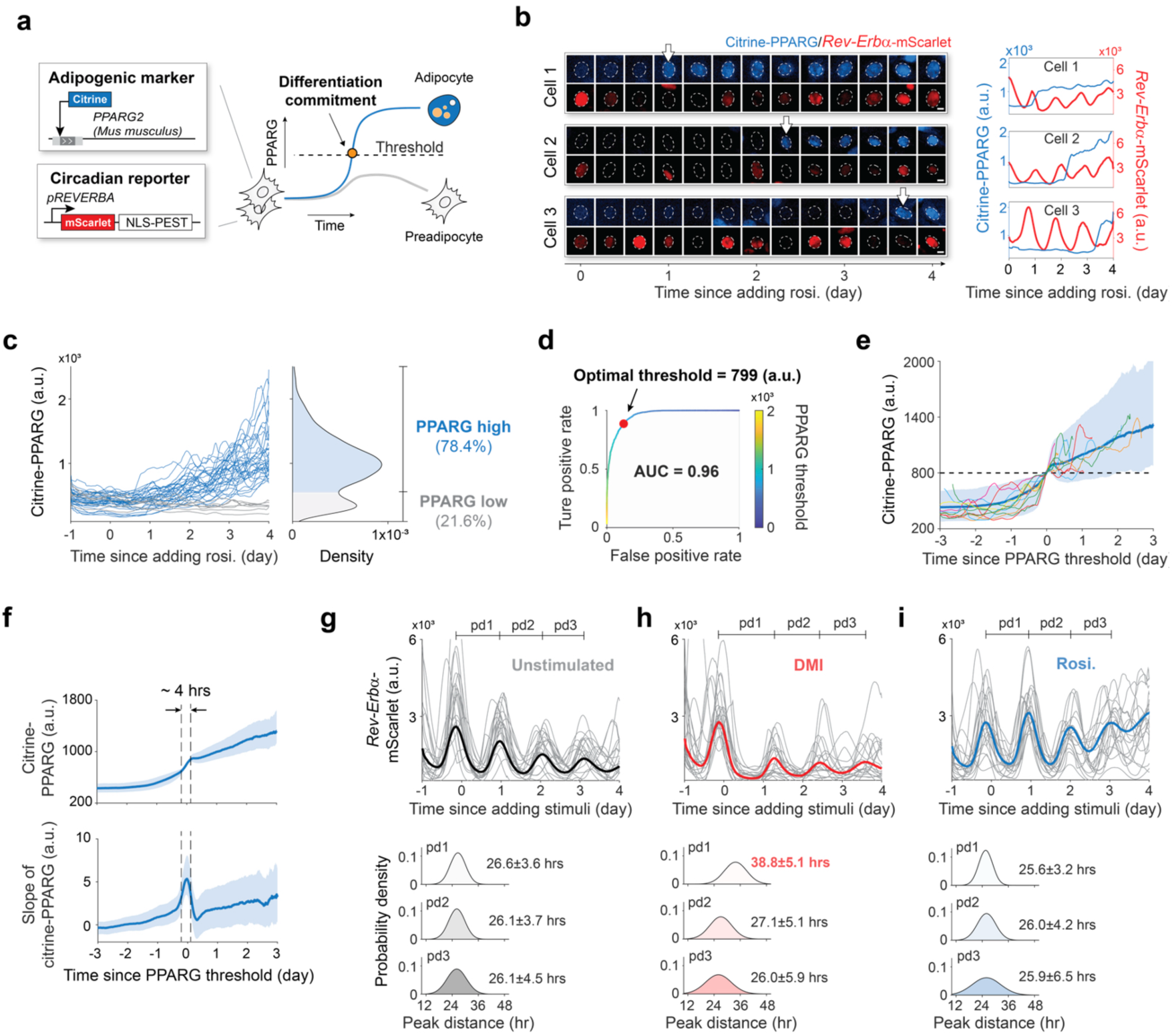
Simultaneously monitoring of the cell-intrinsic circadian clock and adipogenesis in single cells. (**a**) Schematic of the cell model design. (**b**) The citrine-PPARG / *Rev-Erb*a-mScarlet dual-reporter cells were stimulated with 100 nM rosiglitazone (rosi.). The dotted outlines mark the nuclei. The arrows indicate the time when cells switch to the high PPARG state. Scale bar, 5 μm. (**c**) Single-cell time courses of citrine-PPARG were divided into two categories based on the citrine-PPARG intensity at day 4. Representative of 3 biological replicates. (**d**) Receiver operating characteristic curve (ROC) analysis was used to determine an optimal threshold in PPARG level which can predict the fate of most individual cells correctly. AUC represents the two-dimensional area underneath the entire ROC curve. (**e, f**) citrine-PPARG time courses from (**c**) were aligned to the time when the cell reached the optimal threshold. **(e)** Plot of 10 representative aligned time courses (light lines), as well as the median (solid line) and the 5th-95th percentiles (shaded area) from *N* = ∼13,000 differentiated cells. (**f**) Plot of median and the 25th-75th percentiles. (**g** to **i**) Each plot shows 20 representative time courses and the median from *N* = ∼5,000 cells. The peak-to-peak distance (pd) is presented as mean+/-SD. Representative of 3 biological replicates.

Adipogenesis is invariably a bistable process: preadipocytes induced to differentiate end up in either a high or low PPARG state(7, 11), corresponding to being either differentiated or undifferentiated (Fig. 1a,c). Upon addition of a differentiation stimulus, PPARG levels start to increase gradually in preadipocytes(7). However, preadipocytes only irreversibly commit to differentiate when PPARG levels increase to a threshold level at which multiple positive feedbacks to PPARG engage so strongly that PPARG levels stay high, even when the differentiation stimulus is removed(7). The time when cells reach the threshold and irreversibly commit to differentiated can be seen by a step-increase in PPARG level (marked with white arrows in Fig. 1b).

To precisely calculate when cells step-up PPARG and pass the threshold for citrine-PPARG levels, we used receiver operating characteristic (ROC) analysis (Fig. 1d, see Methods)(12). In this analysis, different threshold levels are surveyed to find the one that maximizes the difference between true and false positive rates for predicting cell fate choice(12). For the typical experiment shown in Fig. 1d, the area-under-the curve (AUC) value of 0.96 is close to the maximal value of 1, demonstrating that the optimal citrine-PPARG threshold derived from ROC analysis can be used to measure with high accuracy the precise time when cells commit to differentiate. When the citrine-PPARG traces from Fig. 1c are computationally aligned to the time each cell reached the PPARG threshold, a bimodal switch from low (undifferentiated) to high (differentiated) PPARG levels can be observed (Fig. 1e). As show in in Figs. 1e-f, PPARG increases gradually in preadipocytes induced to differentiate, often over several days before the fast commitment step occurs. After the step increase in PPARG level, PPARG levels do not fall back even if the differentiation stimulus is removed, and PPARG levels often continue to increase gradually for a few more days before cells are fully differentiated. We confirmed that this step increase from a lower to higher PPARG levels occurs rapidly within only a few hours (Fig. 1f).

Using the *Rev-Erb*a-mScarlet reporter to analyze the circadian clock dynamics, we found that the circadian period was approximately 26 hours in unstimulated control cells, (Fig. 1g). Adipogenic stimuli often contain added glucocorticoids which promote differentiation, but also reset the circadian clock in peripheral cells and tissues(13). Indeed, we confirmed that applying the commonly-used DMI stimulus that contains a synthetic glucocorticoid, dexamethasone, perturbs the circadian clock by delaying the first peak of the *Rev-Erb*a reporter by approximately 12 hours after which the peak-to-peak distances return to approximately 26 hours (Fig. 1h). To prevent resetting the clock in the analysis of differentiation, we instead used the PPARG agonist rosiglitazone to induce adipogenesis(14, 15) (Fig. S1), which kept the circadian clock period at approximately 26 hours over the several-day time course of adipogenesis (Fig. 1i).

### Differentiation commitment is almost exclusively triggered during the rising phase of the *Rev-Erb*a reporter

To determine when a cell commits to differentiate relative to the cell-intrinsic clock, we analyzed citrine-PPARG expression and the *Rev-Erb*a reporter simultaneously in the same cells during adipogenesis. A visual inspection of hundreds of single-cell time courses (see representative examples in Fig. 2a) showed great variability in the number of circadian cycles that occur before cells commit to differentiate, ruling out the initial hypothesis that cells delay adipogenesis by counting a fixed number of oscillations. We therefore turned to the second possibility introduced above that the cell-intrinsic circadian clock may trigger commitment at a particular circadian clock phase. To determine the phase when a cell commits to differentiate, we first measured the time when each individual cell reached the PPARG threshold for irreversible commitment. We then used a customized MATLAB script to detect the peaks and troughs in the *Rev-Erb*a reporter oscillations by defining each peak as phase 0 or 2π and each trough as phase π (Fig. 2b, middle). Using a linear fit from peaks to troughs, we then converted the time of differentiation commitment into a circadian phase relative to the cell’s last peak of the *Rev-Erb*a reporter (Fig. 2b, right; Fig. S2a).

**Figure 2.**
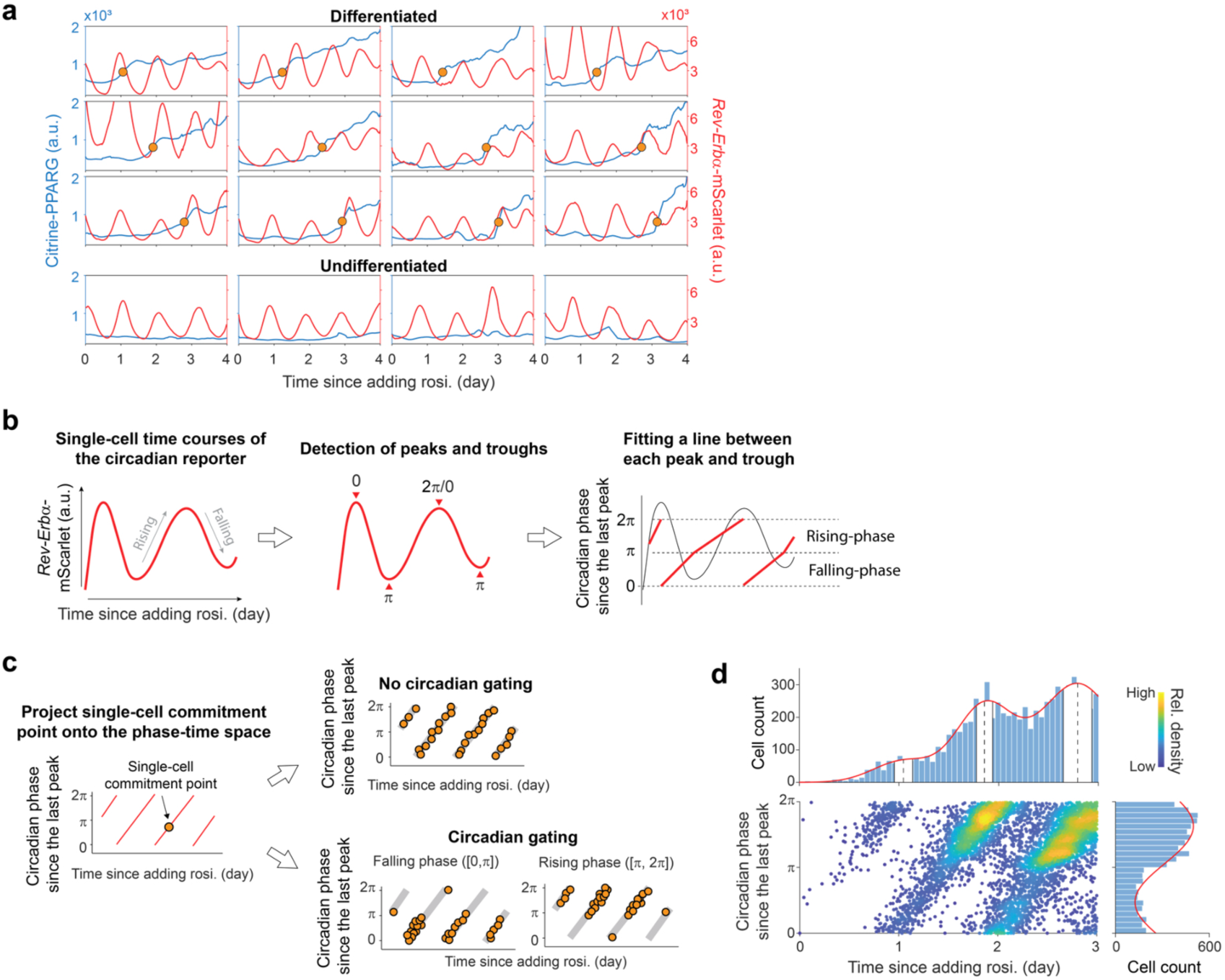
The timing of differentiation commitment follows a circadian pattern. (**a**) Examples of single-cell time courses of citrine-PPARG (blue) and circadian reporter (red) in response to 100 nM rosiglitazone. For differentiated cells, the yellow circle marks the time when the cell reaches the PPARG threshold and commits to the differentiated state. (**b**) Scheme showing procedure to convert the single-cell time courses of the *Rev-Erb*a reporter into the circadian phase. (**c**) Scheme showing how to determine if and how the cell-intrinsic clock regulates the timing of differentiation commitment. (**d**) The scatter plot represents the distribution of commitment points in the phase-time space. *N* = ∼13,000. (top) The distribution of commitment points over time was fitted by a gaussian mixture model. Vertical dashed lines and white bands indicate μ (mean) and σ (SD) for the first three components, respectively. (right) Distribution of commitment point phases was fitted by a one-term Fourier model. Representative of 3 biological replicates.

The scheme in Fig. 2c depicts how this analysis can be used to calculate a phase-corrected time of commitment. The left plot shows the projection of one single-cell commitment point onto the phase-time space. By plotting the distribution of commitment points of thousands of single cells within this phase-time plot, we can determine if and when the cell-intrinsic circadian clock gates differentiation commitment. For example, if a particular time in the circadian oscillation indeed gates when cells commit to differentiate, there should be recurring bursts in the distribution in the phase-time plot during sequential circadian oscillations; otherwise, if cells commit to differentiate independently of the phase of the circadian clock, the commitment points should be evenly spaced over the circadian oscillation (Fig. 2c, right).

We quantitatively tested whether preadipocytes exhibit circadian gating by projecting approximately 13,000 commitment points onto the phase-time plot (Fig. 2d and Fig. S2b). Strikingly, we found that commitment was almost exclusively triggered between the p to 2p half of the circadian cycle, resulting in bursts of differentiation commitment events that were spread over sequential circadian oscillations and provided strong evidence for circadian gating of differentiation. Figure 2d (top) shows that the distribution of the commitment time peaks every day. However, only by also plotting the phase could we also learn that cells preferentially commit to differentiate every day during the rising phase of the *Rev-Erb*a reporter. Since cells were plated at a very high density such that less than 1% could proliferate after being induced to differentiate (Fig. S3), it is unlikely that the non-uniform distribution of the commitment phase is caused by differences in cell cycle phases.

### Manipulations of circadian rhythms demonstrate the cell-intrinsic clock gates the timing of differentiation commitment

Having established that differentiation commitment occurs in bursts that correlate with sequential rising phases of the circadian reporter (Fig. 2d), we next pharmacologically manipulated the circadian oscillation waveforms during differentiation to test whether the rhythms of the circadian clock were indeed controlling the observed gating of differentiation. We first used the commonly used adipogenic cocktail DMI which contains dexamethasone, a potent glucocorticoid that delays the first clock oscillation by approximately 12 hours (Figs. 1g, 3a and Fig. S4a). As shown in Figure 3b, the bursts of differentiation commitment when DMI was added were also shifted by 12 hours. However, even though the differentiation bursts were shifted in time, the phase when cells commit to differentiate remained between p to 2p (Fig. 3b and Fig. S4b), supporting that the rising phase of the *Rev-Erb*a reporter controls differentiation commitment.

**Figure 3.**
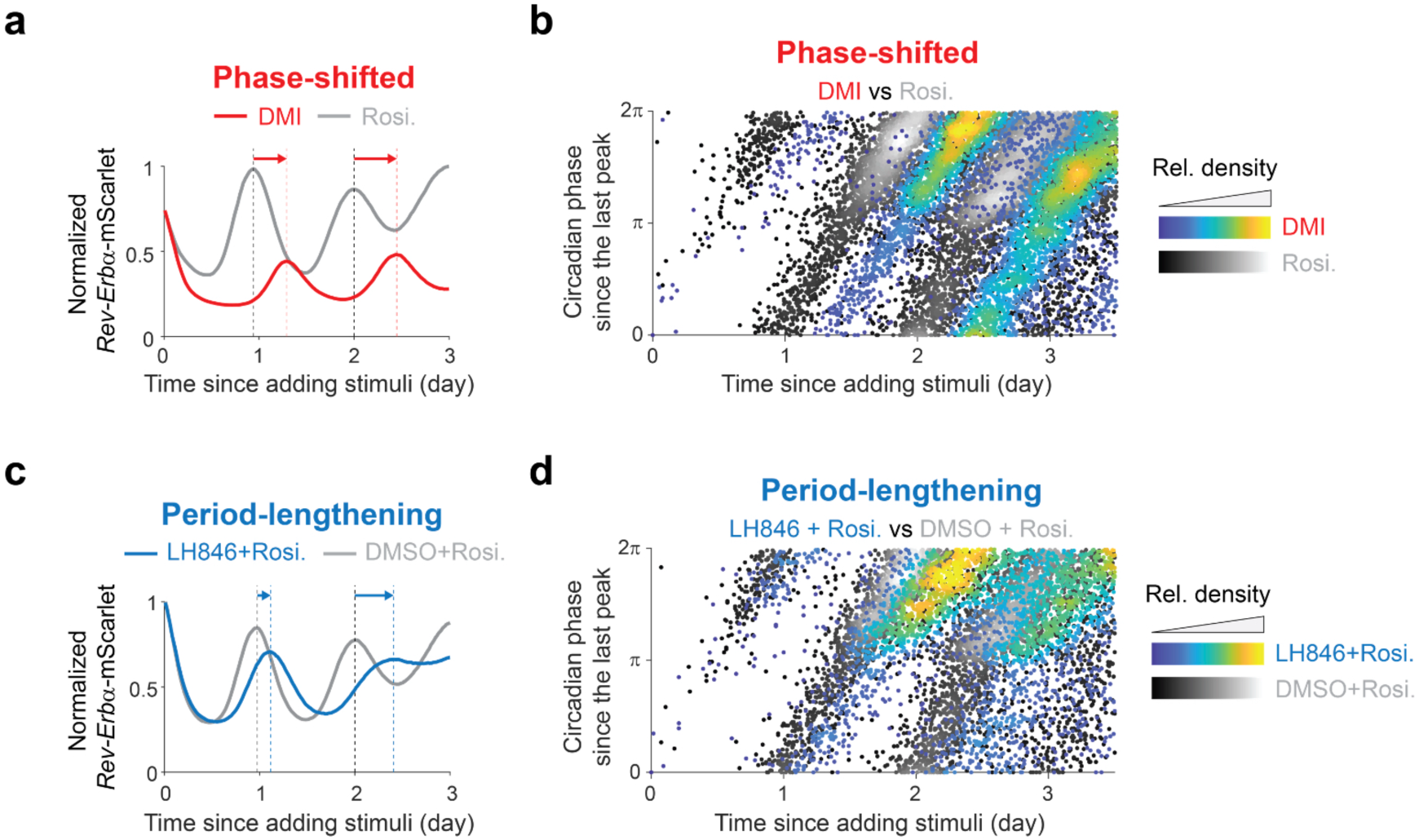
Manipulations of circadian rhythms demonstrate cell-intrinsic clock gates the timing of differentiation commitment. (**a**) The citrine-PPARG / *Rev-Erb*a-mScarlet dual-reporter cells were induced to differentiate by addition of a DMI cocktail or 100 nM rosiglitazone. The median of *N* = ∼5,000 *Rev-Erb*a reporter time courses is normalized to the maximum value. Representative of 3 biological replicates. (**b**) Comparison of the commitment point patterns in the phase-time space shows that differentiation commitment time is pushed back by the delayed phase of the circadian clock, but the time when cells commit to differentiate remains between the π to 2π phase of the circadian reporter (Also see Fig. S4B). (**c**) The citrine-PPARG / *Rev-Erb*a-mScarlet dual-reporter cells were induced to differentiate by 100nM rosiglitazone along with 4 μM LH846 or DMSO (control). The small molecule LH846 caused the periods of the circadian clock to be lengthened. The median of *N* = ∼6,000 *Rev-Erb*a reporter time courses is normalized to the maximum value. Representative of 3 biological replicates. (**d**) The period lengthening caused by the small molecule LH846 resulted in delayed differentiation commitment, but the phases of the commitment points are still enriched in the range from π to 2π (Also see Fig. S4d).

To further validate the role of circadian phase in controlling differentiation commitment, we treated differentiating cells with LH846, a small molecule which lengthens circadian cycles by inhibiting the endogenous degradation of PER proteins(16). Consistent with circadian gating, LH846 gradually delayed the peaks of the *Rev-Erb*a circadian reporter (Fig. 3c and Fig. S4c), but differentiation commitment still occurred tightly between π to 2π of the circadian oscillations (Fig. 3d and Fig. S4d). Taken together, we conclude that differentiation commitment occurs in sequential daily bursts gated by the circadian clock. Since the rising-phase of the *Rev-Erb*a reporter corresponds to the sleep/inactive cycle for both diurnal and nocturnal animals(9, 17–19), our results suggest that preadipocytes commit to differentiate primarily during the evening for humans and during the day for mice.

### Circadian regulation of CEBPA triggers bursts of commitment during adipogenesis

As shown in Figure 1e, differentiation commitment occurs when the abundance of PPARG increases to a threshold level. Thus, in order for differentiation commitment to be gated during the π to 2π phase of the *Rev-Erb*a sensor, PPARG abundance can only be boosted to the threshold only during this 12-hour window in each circadian cycle. Since potential circadian oscillations in PPARG synthesis rate are masked by the gradual overall increase in PPARG abundance during adipogenesis (Fig. 1c, f), we averaged time courses of the *Rev-Erb*a reporter and PPARG abundance, respectively, from about 7,000 cells undergoing adipogenesis. We then plotted the slope of *Rev-Erb*a reporter dynamics and the slope of PPARG abundance versus time to examine how the step increase in the PPARG synthesis rate compares to the clock dynamics (Fig. 4a, left). Markedly, the analysis showed that the synthesis rate of PPARG increases more strongly during the rising phase of the *Rev-Erb*a sensor. When oscillations in the circadian clock are abolished by knocking down the key clock protein BMAL1, PPARG synthesis still increases overall in response to the differentiation stimulus, but not in an oscillatory fashion (Fig. 4a, right).

**Figure 4.**
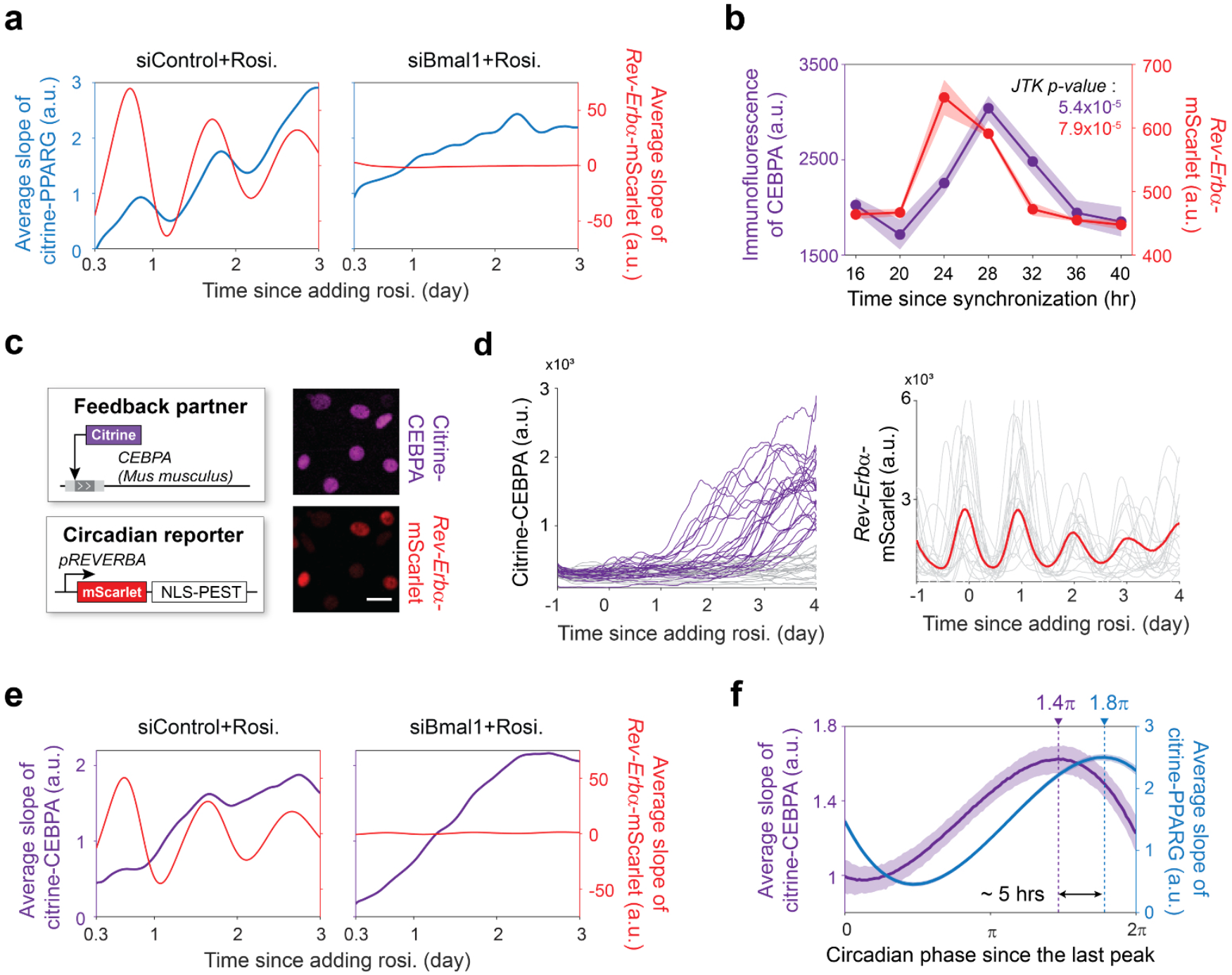
Both PPARG and its feedback partner CEBPA are expressed in a circadian pattern during adipogenesis. (**a**) The average slope of PPARG time courses from *N* = ∼7,000 cells shows that the PPARG synthesis rate follows a circadian pattern. Representative of 3 biological replicates. (**b**) Immunofluorescent staining was performed for CEBPA after cells were synchronized by a 1-hour dexamethasone pulse. Bar plot represents mean+/-SD of 3 technical repeats with *N* = ∼10,000 cells each. (**c**) System to simultaneously monitor the dynamics of CEBPA and circadian rhythm. Scale bar, 10 μm. (**d**) The citrine-CEBPA / *Rev-Erb*a-mScarlet dual-readout cells were induced to differentiate using 100 nM rosiglitazone. *Left*, single-cell time courses of citrine-CEBPA can be divided into two categories (purple, grey) based on the nuclear citrine-CEBPA intensity at day 4. *Right*, plot shows 20 representative time courses and the median from *N* = ∼5,000 cells. (**e**) The average slope of CEBPA time courses of *N* = ∼5,000 cells shows that CEBPA synthesis rate also follows a circadian pattern. Representative of 3 biological replicates. (**f**) Single-cell time courses were computationally aligned and scaled to circadian phase from 0 to 2π for each period. Plotted are the mean and the 95% confidence intervals generated from 1,000 bootstrap resampling at each timepoint. Dashed lines indicate the circadian phase of the maximum slope values.

What causes PPARG synthesis rate to oscillate in a circadian fashion is puzzling since PPARG does not have BMAL1/CLOCK-regulated E-boxes in its promoter. E-boxes are typically needed to control the circadian expression of genes. Since PPARG is regulated by several positive feedback regulators(7, 20), we considered that one of these partners may instead be regulated by BMAL1/CLOCK. We focused on CEBPA since it is the main positive feedback partner of PPARG(21), has two E-boxes on its promoter(22), and its mRNA level was shown to oscillate in a circadian pattern in fibroblasts(22). We first knocked down CEBPA and assessed the effect on PPARG circadian oscillations. Knockdown of CEBPA dramatically reduced the expression of PPARG, as well as the amplitude of the slope of the circadian oscillations (Fig. S5), supporting that CEBPA could be driving PPARG circadian oscillations.

To test whether CEBPA expression is circadian in OP9 cells, we plated OP9 cells in 96-well plates and added to each well a 1-hour pulse of dexamethasone, a synthetic glucocorticoid which has been shown to synchronize the circadian clock(10). The pulse duration of dexamethasone was selected to be too short to induce differentiation. We then collected cells every 4 hours from different wells and carried out immunofluorescence analysis to track protein abundance. Consistent with a circadian regulation, CEBPA expression followed a circadian pattern based on a JTK_Cycle rhythmicity test(23) and peaked shortly after the peak in *Rev-Erb*a reporter expression (Fig. 4b). To understand the relationship between circadian expression of CEBPA and circadian PPARG expression, we used CRISPR-mediated genome editing to tag endogenous CEBPA with citrine (YFP) in OP9 preadipocyte cells (Fig. 4c). By stably transfecting the *Rev-Erb*a-mScarlet reporter into these cells, we could monitor CEBPA activity and circadian rhythms simultaneously in the same cell (Fig. 4d). We induced adipogenesis of the dual reporter citrine-CEBPA / *Rev-Erb*a-mScarlet cells and calculated how the slope of the citrine-CEBPA level changes over time relative to the circadian cycles. Indeed, performing the same slope analysis, as done in Figure 4a for PPARG, confirmed that CEBPA synthesis rate increases in a circadian manner (Fig. 4e, left). However, when Bmal1 is knocked down, the rate of CEBPA synthesis no longer increases in an oscillatory manner although it does increase overall in response to the adipogenic stimulus (Fig. 4e, right).

To more precisely quantify when the rate of CEBPA protein synthesis increases, we measured the change in the citrine-CEPBA signal during each circadian period (measured by the peak-to-peak distance in the *Rev-ERBa-*mScarlet signal, 0 to 2 π in Fig. 2b) for the cells in Fig. 2d to obtain an average citrine-CEBPA slope during a circadian period (Fig. 2F). We found that the citrine-CEBPA slope took the shape of a sine wave peaking at ∼1.4π (Fig. 4f). In contrast, the circadian oscillations of PPARG peaked slightly later at ∼1.8π. Because the average circadian period is about 26 hours (Fig. 1g), the gap between the peaks of CEBPA and PPARG slopes correspond to an approximate 5-hour delay (Fig. 4f). Together, these results support clock-induced expression of CEBPA during each rising phase of the *Rev-Erba* reporter drives circadian PPARG expression.

Our experiments in Fig. 4 showed that clock-driven peaks of CEBPA expression are correlated with with the peaks of PPARG expression. To now determine whether clock-driven CEBPA is what drives the daily bursts of differentiation commitments, we used our previous computational model to simulate PPARG dynamics in response to an adipogenic stimulus(7, 11). Our model includes that differentiation commitment during adipogenesis is driven by fast and slow positive feedbacks centered on PPARG(7, 20) (Fig. 5a). We used t_1/2_ = 3 hours for the fast regulation which is the timescale of the CEBPA-PPARG feedback loop and a t_1/2_ = 30 hours for the slow regulation which is the timescale of the FABP4-PPARG feedback loop(7). As shown in Figure 5b, the simulations recapitulate our experiments which showed that the overall PPARG level in cells increases slowly over days, but close to the threshold, there is a more rapid increase in PPARG level as cells irreversibly commit to differentiate (Figs. 1c, 1e, and 1f). Furthermore, the simulations recapitulate known cell-to-cell variability (noise) in the fast and slow feedback circuits that regulate PPARG (7, 20, 24): cells reach the threshold at different times after adipogenesis is induced, but in a manner that is evenly spaced over a few days, as is readily apparent from the histogram in Fig. 5b. Markedly, when we added a term to the model that superimposes oscillating circadian synthesis of CEBPA (Fig. 5c), we recapitulated the experimentally observed daily bursting behavior of cell differentiation (Fig. 2d). Thus, our simulations support that clock-driven CEBPA-expression can drive circadian PPARG expression to generate daily bursts of cell differentiation over multiple clock cycles.

**Figure 5.**
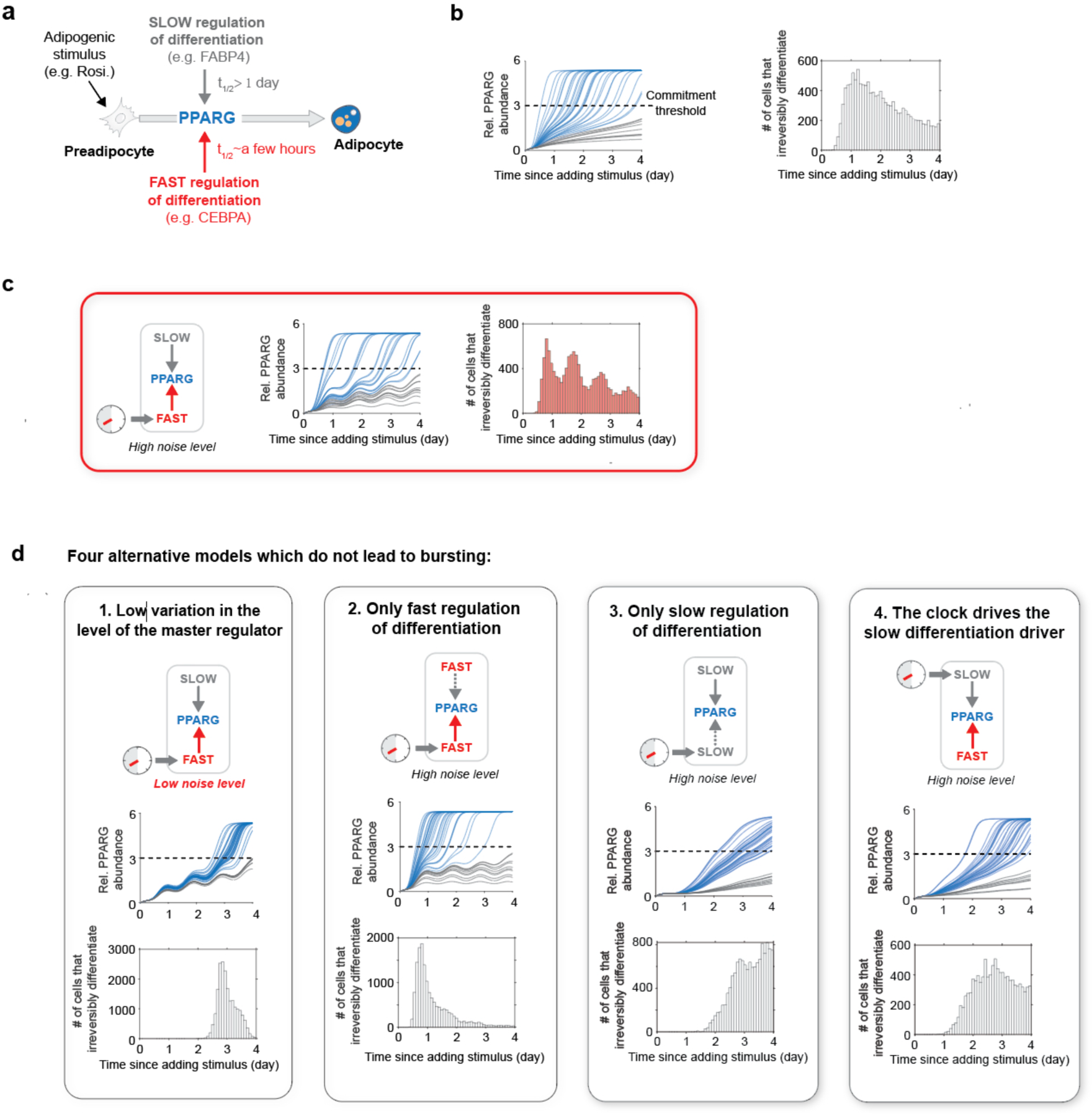
Four requirements for circadian gating of differentiation commitment and generating daily bursts of cell differentiation. (**a**) Schematic of canonical regulatory circuits controlling the expression of PPARG during adipogenesis. (**b**) Quantitative simulations show that due to the combined fast and slow feedback regulation, the abundance of PPARG increases slowly before and after a rapid switch that occurs as cells reach the threshold (dashed line). 20,000 simulations were carried out, and the cell-to-cell variability was taken into account by randomly adding 30% log-normal noise to each simulation. Blue lines represent representative differentiated cells whose PPARG level passed the threshold line, with the gray lines representing undifferentiated cells. As consistent with the live-cell analysis, the time of differentiation commitment was defined as the time when PPARG level reached the threshold. (**c**) Coupling the circadian clock to the fast regulator CEBPA in the model recapitulates the experimentally-observed circadian bursts of cell differentiation. (**d**) Four simulations in which a different regulatory element was changed. High variation, slow and fast regulation of the master regulator, and coupling of the clock to the fast regulator are needed to generate daily bursts of differentiation commitments.

## Discussion

Our experiments and modeling showed that cell differentiation is gated, meaning that the circadian clock restricts differentiation commitment of individual preadipocytes almost exclusively to the rest phase each day. We now discuss how circadian gating of individual cells can lead to the bursting behavior at the level of the cell population as seen in Figs. 2d and 5c.

In the individual timecourses of citrine-PPARG and *Rev-Erba*-mScarlet dual reporter cells (Fig. 2a), we can see that there is variability when a cell irreversibly decides to commit to differentiate. In each clock cycle after differentiation is induced, only a subset of the progenitor cells reach the PPARG threshold to irreversibly commit. Because only some individual cells commit to differentiate in each of several consecutive circadian cycles, a bursting behavior can be observed at the population level. The reason only some cells reach the threshold at a given time is likely because of the previously described cell-to-cell variability in the expression level of the master regulator PPARG when adipogenesis is induced(7, 20, 24). To validate the importance of cell-to-cell variation in PPARG expression for generating the bursting behavior, we reduced the variation of PPARG in the model. As shown in the traces and the single peak in the histogram in Fig. 5d.1, reducing cell-to-cell variation of PPARG expression indeed leads to a loss of the bursting behavior.

In the model that generated bursting (Fig. 5c), we had included fast and slow regulation of the master differentiation regulator. We now wanted to understand if both speeds of regulation are needed to generate multiple daily bursts of cell differentiation. We thus modified the model to have only fast or only slow regulation of PPARG. As shown in the histogram in Fig. 5d.2, with only fast regulators, almost all cells differentiate in the first clock cycle. As shown in the histogram in Fig. 5d.3, with only slow regulators, cells commit to differentiate over a broader range of clock cycles. However, PPARG levels do not behave in a circadian manner since the slow regulation prevents rapid degradation of PPARG(7, 25), and thus, PPARG levels cannot rapidly drop during the waking phase of the circadian clock. These results support that both slow and fast control of the master differentiation regulator are needed to generate circadian differentiation bursts over multiple days. We also wanted to understand whether the circadian clock must drive the fast regulator of differentiation. As shown in Fig. 5d.4, when we modified the model to have the clock to drive the slow regulator, no circadian expression of PPARG was observed and cells did not commit to differentiate in a circadian manner.

Our results support that there are 4 requirements to generate circadian differentiation bursts over multiple days: (1) high cell-to-cell variability in expression of the master differentiation regulator; (2) a slow differentiation driver; (3) a fast differentiation driver, and (4) the circadian clock must be coupled to the fast differentiation driver. Cell-to-cell variability and slow regulation of PPARG prevent cells from all differentiating at one time or immediately after differentiation is induced. Fast regulation of PPARG allows differentiation commitment to occur rapidly – in approximately 4 hours (Fig. 1f) – which is well within the duration of the rest phase of the circadian clock. Coupling the clock to CEBPA, which is in a fast feedback relationship with PPARG, allows PPARG levels in individual cells to be boosted rapidly towards the threshold during each rest phase of the clock. If an individual cell does not reach the threshold to differentiate within this gating window, the fast regulation of PPARG allows PPARG levels to quickly drop back down during the active phase of the clock, generating a circadian pattern of PPARG expression. Thus, coupling fast regulation of PPARG to the circadian clock explains the gating of differentiation commitment in individual cells to the rest phase of the circadian clock, and cell-to-cell variability and the slow regulation of PPARG explain why differentiation commitment is spread out over multiple clock cycles and can thus explain the bursting behavior at the population level.

Overall, our study provides direct evidence that the circadian clock restricts fat cell differentiation commitment to the rest phase each day. Clock-mediated restriction of the differentiation commitment step, which involves major transcriptional and chromatin changes, to the rest phase, during which metabolic activity is likely lower than in the wake period, may help to increase the reliability of cell differentiation. Our study also defines the differentiation system criteria needed to generate the observed daily bursts of cell differentiation. Other cell differentiation systems such as Th17 and skeletal muscle differentiation may employ similar circadian bursting and gating regulation mechanisms (4, 26).

## Materials and Methods

### Generation of citrine-PPARG / *Rev-Erbα*-mScarlet dual readout cell lines

To generate a stable live cell sensor for *Rev-Erbα* activity, the entire open reading frame of the Rev-VNP expression cassette described in a previous work(10) was PCR amplified in addition to a 1kb region upstream of the start codon containing the *Rev-Erbα* promoter elements. The amplified fragments were cloned using Gibson assembly into a Piggyback expression backbone PB-CMV-MCS-EF1α-Puro vector (System Biosciences), which has been previously modified with PGK-Blasticidin in place of pEF1α-Puromycin and linearized using SfiI/XbaI. The assembled construct, PB-REVERBA-Venus-NLS-PEST, was then digested with NotI and SalI to swap Venus fluorophore with a GBlock-Gene Fragment (IDT) containing mScarlet which was inserted using Gibson assembly to generate PB-REVERBA-mScarlet-NLS-PEST. The PB-REVERBA-mScarlet-NLS-PEST construct was then transfected into OP9 cells already expressing endogenously tagged citrine(YFP)-PPARG(7) using Lipofectamine 2000 (Thermofisher). Cells were selected for 48 hours post transfection using 10 μg/ml Blasticidin (Invivogen) for 10 days and FACS sorted for mScarlet (RFP). To facilitate cell tracking in microscopy experiments, cells were subsequently infected with lentivirus (PLV-H2B-mTurquoise) to introduce a nuclear marker and further FACS sorted on CFP. During this process, single clones were also isolated and expanded and tested for their ability to maintain proper circadian rhythmicity and differentiate into adipocytes upon DMI stimulation.

### Generation citrine-CEBPA / *Rev-Erbα*-mScarlet dual readout cell lines

To generate OP9 cells in which endogenous CEBPA is tagged at the N-terminus with citrine, we followed the same protocol used to tag the N-terminus of endogenous PPARG in OP9 cells with citrine (YFP)(7). The nuclear marker (PLV-H2B-mTurquoise) and circadian reporter (PB-REVERBA-mScarlet-NLS-PEST) were then stably integrated into the citrine-CEBPA cells. Single clones were isolated and tested in the same manner as described above.

### Cell culture and differentiation

The wildtype OP9 cells and the dual readout OP9 cells were maintained according to previously published protocols(7). Briefly, the cells were cultured in full growth media consisting of MEM-α media (ThermoFisher Scientific) containing 1 unit/mL Penicillin, 1 mg/mL Streptomycin, and 292 μg/mL L-glutamate supplemented with 20% FBS. To induce differentiation, 100 nM of rosiglitazone (Cayman, USA) or the adipogenic cocktail (DMI) consisting of dexamethasone (1 μM, Sigma-Aldrich), IBMX (250 μM, Sigma-Aldrich), and insulin (1.75 nM, Sigma-Aldrich) were used. For live-imaging experiments, the differentiation stimuli were added to Fluorobrite DMEM media (ThermoFisher Scientific) supplemented with 10% FBS, and then cells would be continually imaged for 4 days. The small molecule LH846 (Cayman, USA) was used as a concentration of 4 μM. For fixed-cell experiments, stimuli were added to MEM-α media (ThermoFisher Scientific) supplemented with 10% FBS for 2 days, and then removed and replaced with fresh media containing 1.75 nM insulin (Sigma-Aldrich) and 10% FBS for another 2 days.

### Live-cell imaging

15,000 cells per well were plated 24 hours prior to imaging in full growth media in Ibidi μ-Plate (#89626). Before image acquisition, the full growth media was aspirated and replaced with fresh Fluorobrite DMEM media (ThermoFisher Scientific) supplemented with 10% FBS to reduce background fluorescence. Live-cell imaging was conducted using the ECLIPSE Ti2 inverted microscope (Nikon) with a 10X Plan Apo 0.45 NA objective. Cells were imaged in a humidified 37°C chamber at 5% CO2, and images were taken every 12 min in 3 fluorescent channels: CFP, YFP and RFP. Total light exposure time was kept to less than 600 ms for each time point. Four non-overlapping sites in each well were imaged.

### Immunofluorescence staining and imaging

Cells were plated in Costar 96-well plates (#3904) and fixed with 4% PFA in PBS for 15 min at room temperature, followed by four washes with PBS using an automated plate washer (Biotek). Cells were then permeabilized with 0.1% Triton X-100 in PBS for 20 minutes at 4°C, blocked with 5% bovine serum albumin (BSA, Sigma Aldrich) in PBS for 1 hour at room temperature and stained with primary antibody (rabbit anti-CEBPA, 1:1000, Santa Cruz Biotech #sc-61; rabbit anti-PPARG, 1:1000, Cell Signaling #2442; mouse anti-PPARG, 1:1000, Santa Cruz Biotech #sc-7273; Goat anti-FABP4, 1:1000, R&D Systems #AF1443; Mouse anti-Adiponectin, 1:1000, Abcam #ab22554; Goat anti-Glut4, 1:1000, Santa Cruz Biotech #sc-1608) in 1% BSA overnight at 4°C. After four washes, cells were incubated with Hoechst (1:2000) and secondary antibody (Alexa Fluor 647 anti-rabbit, 1:1000) in dark in 1% BSA for 1hour at room temperature. Prior to imaging, cells were washed four times with PBS. For assays involving EdU staining, cells were treated with 10 μM EdU for about 15 min prior to fixation. Fixed-cell imaging was conducted using an ImageXpress MicroXL automated epifluorescence microscope (Molecular Devices, USA) with a 10X Plan Apo 0.45 NA objective. Several non-overlapping sites in each well were imaged.

### Image processing and analysis

Fluorescence images were analyzed using custom scripts and the MACKtrack package(8) in MATLAB R2021a (MathWorks). Cells were segmented and tracked for their nuclei based on either Hoechst staining (fixed cell imaging) or H2B-Turquoise marker (live cell imaging). Flat-field correction for each channel was carried out prior to signal measurement. Quantification of PPARG, CEBPA, mScarlet and EdU in cells was based on quantification of mean fluorescence signal over nuclei. Each single-cell trajectory was smoothed using a moving average filter with a 6-hour span. The slope of PPARG, CEBPA and *Rev-Erbα*-mScarlet at each timepoint was calculated by using a linear fit to 8-hour segments of the trajectory (+/-4 hours).

### Calculating the threshold for differentiation commitment

The terminal fate for a given cell was scored as differentiated or undifferentiated based on if its terminal PPARG expression level was above or below a preset cut-off value. The preset cut-off value (the ground truth) was set so that there will be less than 3% of control (DMSO) cells scored as terminally differentiated cells (Fig. S6). Then, to determine when cells commit to the differentiated state, we tested a series of thresholds to predict cells’ terminal fates before the experiment ended. For a given threshold value, cells would be predicted as differentiated if their nuclear citrine-PPARG time courses reached above the threshold value prior to the end of the experiment. The false positive rate and true positive rate of the predictions were calculated based on the ground truth. Next, we plotted the receiver operating characteristic curve (ROC) for all the threshold values and selected the one as the optimal threshold whose point on the ROC was closest to the corner point (0,1). This optimal threshold can maximize the difference between true positive rate and false positive rate for predicting cell fate choice. The time of differentiation commitment for each terminally differentiated cell was determined as the moment when its nuclear citrine-PPARG time course crossed the optimal threshold for the first time.

### siRNA-mediated gene silencing

siRNA targeting Bmal1 and the AllStars Negative Control siRNA were purchased from Qiagen. For siRNA knockdown in the live-cell imaging experiments, dual readout cells were transfected by reverse-transfection using Lipofectamine RNAiMax (Invitrogen). Briefly, our reverse-transfection protocol per well is as follows: mix 40 μL of Opti-MEM media (ThermoFisher Scientific), 0.4 μL of a 10 μM siRNA stock solution, and 1 μL of RNAiMax. The solution was incubated at room temperature for 10 minutes, and then 160 μL of culture media containing the desired number of cells per well was added. Then the entire (∼200 μL) volume was plated into one well of an Ibidi 96-well μ-plate. The siRNA/RNAiMax mixture was left on the cells for 24 hours before being aspirated away and replaced with fresh Fluorobrite DMEM media supplemented with 10% FBS prior to imaging.

### Statistics

Statistical parameters are reported in the figures and figure legends. All statistical analysis was performed in MATLAB R2021a (MathWorks) or R.

### Mathematical modeling

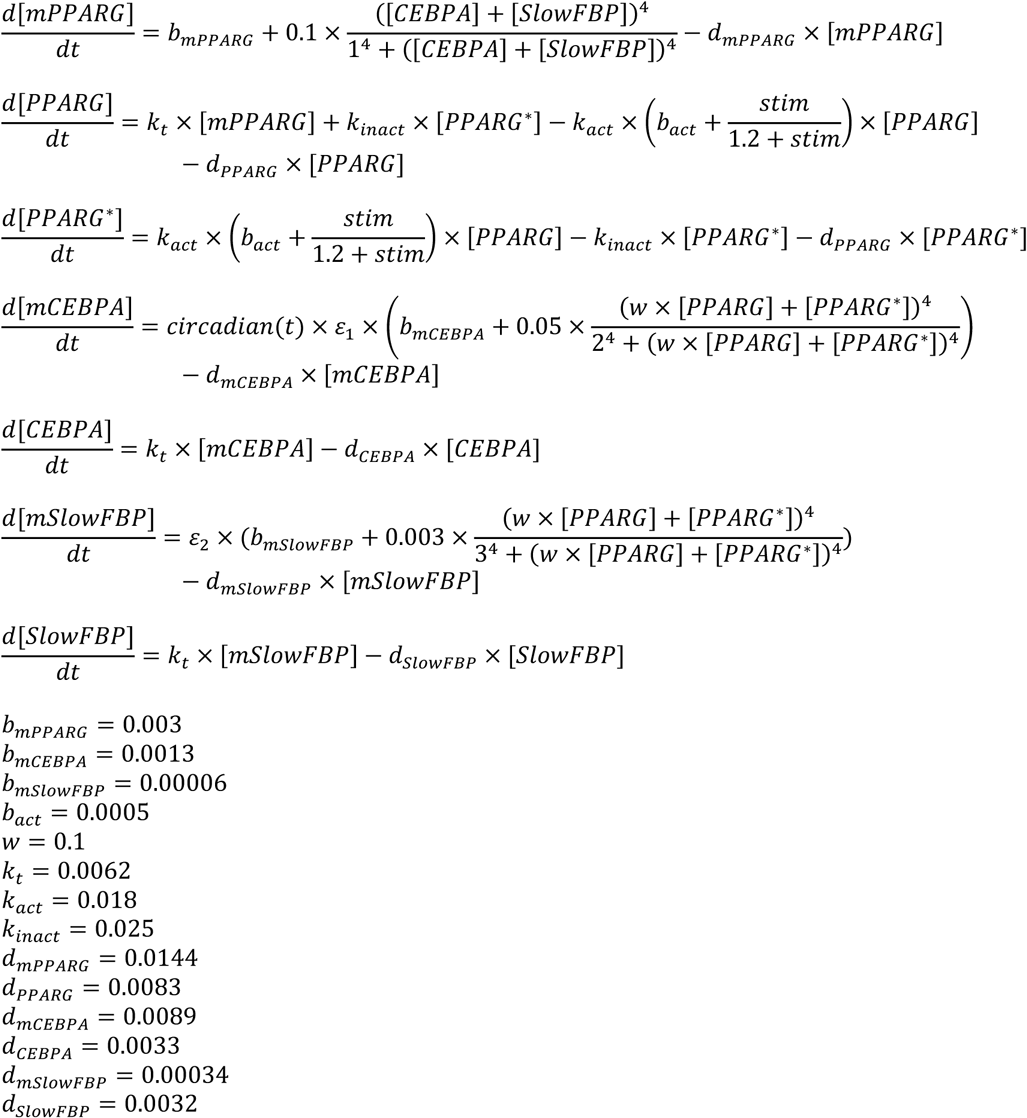

1. The rate of change of the concentrations of seven species are calculated: PPARG mRNA, inactivated PPARG protein, activated PPARG protein (*PPARG*\*), CEBPA mRNA, CEBPA protein, slow feedback partner mRNA (*mSlowFBP*) and slow feedback partner protein (*SlowFBP*).
2. All the variables are initialized to be zero.
3. *k_t_* represents translation rate.
4. The factor w represents the relative activity of the original PPARG and the agonist activated PPARG.
5. Degradation rates are adopted from previous measurements(7).
6. To mimic the adipogenic stimulus, stim is set to be 1 at day0.
7. A cell is scored as differentiated if the concentration of total PPARG protein ([*PPARG*]+[*PPARG*\*]) is above a cut-off determined by the bimodal expression at the end of the simulation.
8. The term *circadian(t)*, in the equation describing CEBPA transcription rate, is set to be a time-dependent cosine function.
9. In the scenario of slow-slow architecture (Fig. 5d), the degradation rates of CEBPA mRNA and CEBPA protein were replaced with that of slow feedback partner mRNA (*mSlowFBP*) and slow feedback partner protein (*SlowFBP*).
10. Lognormal noise (with mean = 0, standard deviation = 30% for high noise level and standard deviation = 3% for low noise level) is randomly added to simulations through three independent noise factors (ε1 and ε2) before the synthesis terms of CEBPA mRNA and slow feedback partner mRNA(7, 20).

## Data availability

Data and custom codes used for the analysis presented in the paper are available at https://hdl.handle.net/1813/110377. Raw data are available from the corresponding author upon request.

## Acknowledgements

We thank Ueli Schibler (University of Geneva) for kindly sharing the Rev-VNP plasmid. We thank Tobias Meyer (Weill Cornell Medicine) and members of the Teruel lab and the Meyer lab for helpful discussions and critical reading of the manuscript. This work was supported by National Institutes of Health RO1-DK101743, RO1-DK106241, R56-DK131423, and a Stanford Diabetes Research Center Seed Grant (to M.N.T.) and T32-NIH GM113854 (to J.S.).

## Notes

### Competing Interest Statement

The authors have declared no competing interest.

