## Supplemental Information for "The circadian clock mediates daily bursts of cell differentiation by periodically restricting cell differentiation commitment"

### The cell-intrinsic circadian clock gates fat cell differentiation commitment

**This PDF file includes:**

Figures S1 to S6

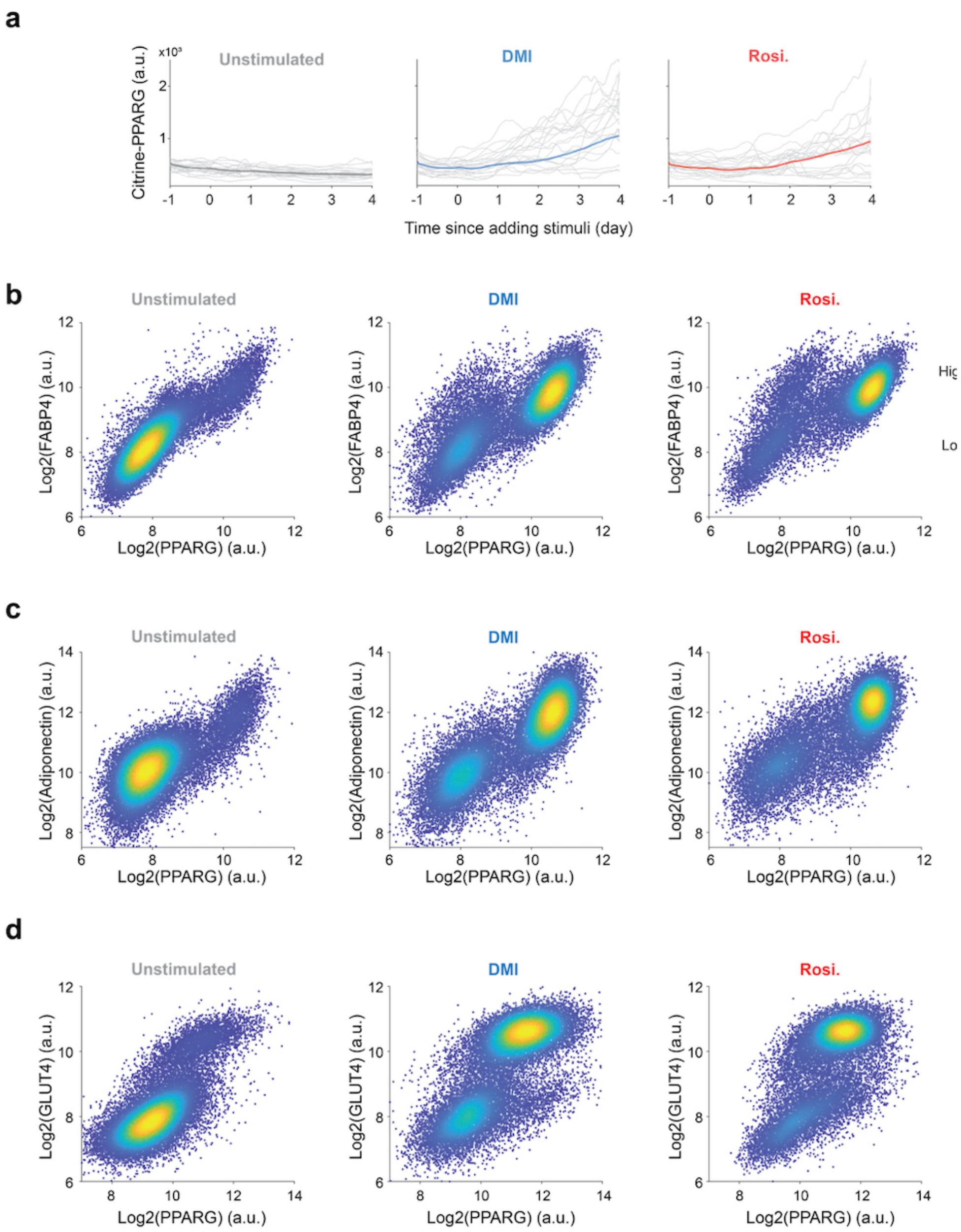


**Fig. S1 Rosiglitazone can induce OP9 preadipocyte cells to differentiate into fat cells.**

**a,** The citrine-PPARG / *Rev-Erb*$\alpha$-mScarlet dual-readout cells were stimulated by DMSO (Unstimulated), adipogenic cocktail DMI or 100 nM rosiglitazone at day 0. Time courses of nuclear citrine-PPARG show that both rosiglitazone and DMI can induce the expression of PPARG in OP9 preadipocyte cells. The bold lines represent population median of N ~17,000 cells and the fine lines represent 20 single-cell time courses for each condition. **b, c, d,** Wild type OP9 preadiopcyte cells were stimulated by DMSO (Unstimulated), adipogenic cocktail DMI or 100 nM rosiglitazone for 48 hours. The media were replaced with fresh media just containing insulin for another 48 hours. Immunocytochemistry for adipocyte markers (**b**. FABP4, **c**. Adiponectin, **d**. GLUT4 along with PPARG) was performed at 96 hour. Like the commonely-used adipogenic cocktail DMI, 100 nM rosiglitazone can also induce the expression of adipocyte markers in OP9 cells.

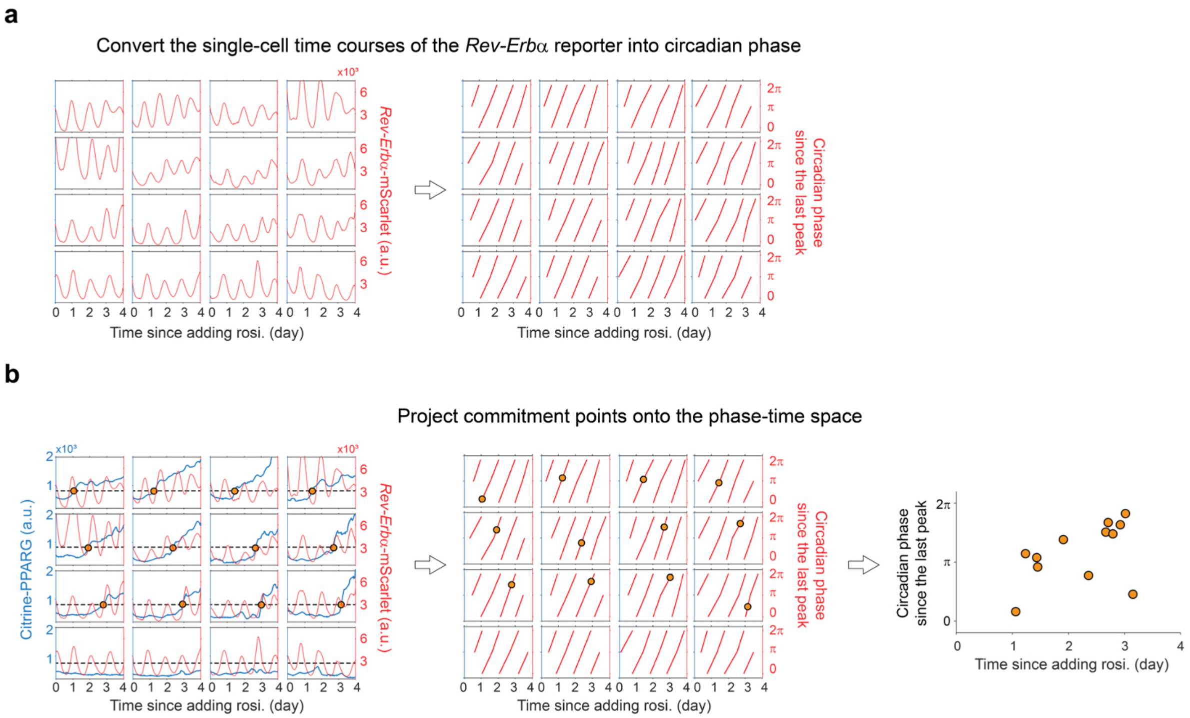


**Fig. S2 Demonstrating how to project the commitment points of differentiated cells into the phase-time space.**

**a,** The *Rev-Erbα*-mScarlet time courses of 16 representative cells from Figure 2a were converted into circadian phase time courses according to the procedures in Figure 2b. **b,** The optimal threshold for PPARG level (horizontal dashed lines) was used to determine if and when the cell commits to the differentiated state. The time of the commitment point for a terminally differentiated cell was converted into circadian phase according to its circadian phase time courses calculated in **b**. The commitment points of all differentiated cells were then projected onto the phase-time space.

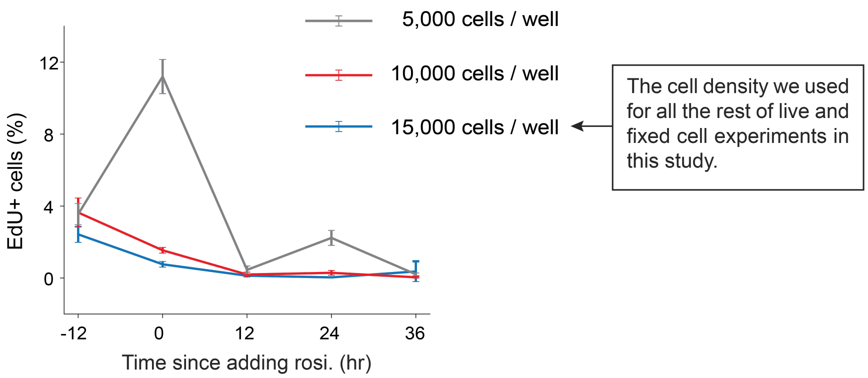


**Fig. S3 Cells were plated at a high density to reduce the percentage of proliferating cells.**

The citrine-PPARG / *Rev-Erb*$\alpha$-mScarlet dual-readout cells were plated at three different densities and induced to differentiate using 100nM rosiglitazone. Cells were stained with EdU and fixed at multiple time points. The percent of EdU+ cells shows mean+/-SD from 3 technical replicates. Representative of two biological replicates.

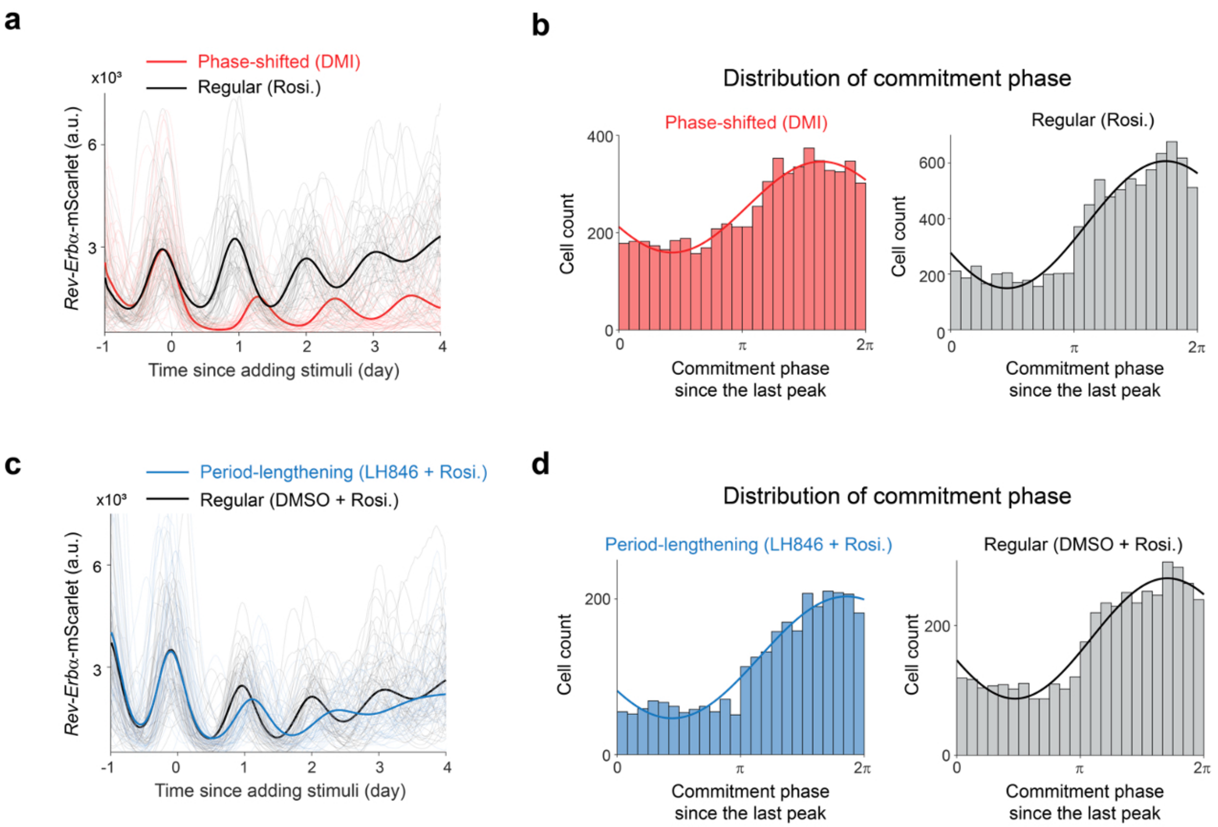


**Fig. S4 Shifted phases or lengthened periods do not affect the preferentially differentiation commitment of preadipocytes in the rising phase of the *Rev-Erbα* reporter.**

**a,** The citrine-PPARG / *Rev-Erbα*-mScarlet dual-readout cells were stimulated by DMI cocktail or 100 nM rosiglitazone at day 0. Time courses of nuclear mScarlet show that circadian clock is delayed by DMI. The bold lines represent population median of ~ 17,000 cells and the fine lines represent 50 single-cell traces for each condition. Representative of 3 biological replicates. **b,** The distribution of commitment phase for the differentiated cells in **a** was fitted by a one-term Fourier model. **c,** The citrine-PPARG / *Rev-Erbα*-mScarlet dual-readout cells were stimulated by 100 nM rosiglitazone along with 4 μM LH846 or DMSO at day 0. Time courses of nuclear mScarlet show that circadian clock is slowed down by LH846. The bold lines represent population median of ~ 7,000 cells and the fine lines represent 50 single-cell traces for each condition. Representative of 3 biological replicates. **d,** The distribution of commitment phase for the differentiated cells in **c** was fitted by a one-term Fourier model.

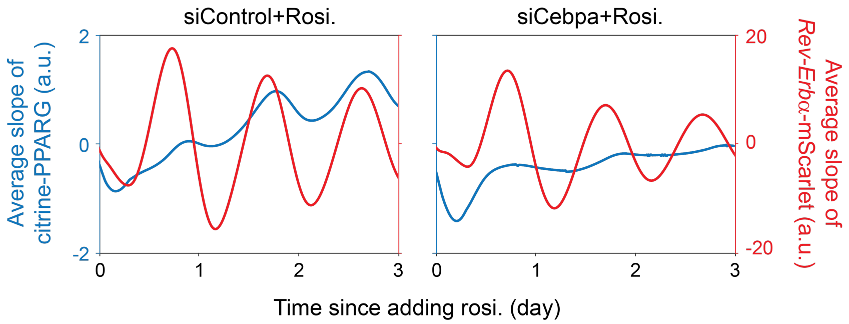


**Fig. S5 Knockdown of CEBPA reduced the amplitude of the circadian oscillations of PPARG synthesis rate.**

The average slope of PPARG time courses from N = ~8,000 citrine-PPARG / *Rev-Erb*$\alpha$-mScarlet dual-readout cells shows that knockdown of CEBPA reduced the amplitude of the circadian oscillations of PPARG synthesis rate.

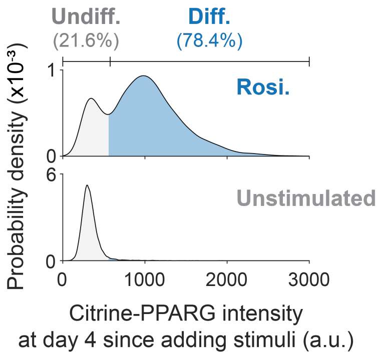


**Fig. S6 Determine cell terminal fate based on the PPARG distribution after adding stimulus for 4 days.**

The citrine-PPARG / *Rev-Erb*$\alpha$-mScarlet dual-readout cells were stimulated by DMSO (Control) or 100 nM rosiglitazone at day 0. The terminal fate for a given cell was scored as differentiated (diff.) or undifferentiated (undiff.) based on if its PPARG expression level at day 4 was above or below a preset cut-off value. The preset cut-off value was set so that there will be less than 3% of unstimulated (DMSO treated) cells scored as terminally differentiated cells.
